# Role of Dengue in SARS-CoV-2 Evolution in Dengue Endemic Regions

**DOI:** 10.1101/2025.04.13.648564

**Authors:** Abinash Mallick, Subhajit Biswas

**Author notes:** Correspondence: Subhajit Biswas, Infectious Diseases and Immunology Division, CSIR-Indian Institute of Chemical Biology, Kolkata, West Bengal, India. **Email:**.

## Abstract

Evidence suggests that dengue virus (DV) antibodies (Abs) and SARS-CoV-2 Abs were cross-reactive, resulting in reciprocal serological cross-interaction and providing cross-protection in populations in co-endemic regions. It became apparent from the present study that SARS-CoV-2 variants were preferentially selecting mutation(s)/deletions in the spike to evade interaction with DV Abs.

We looked at mutations in the SARS-CoV-2 spike protein and how they affected the antigenicity, focusing on successively emergent major variants such as B.1.1.7 (Kent), B.1.617 (Delta), B.1.617.2.1 (Delta-Plus), B.1.1.529 (Omicron-BA.1) and B.1.1.529 (Omicron-BA.2). To further understand the effect of cross-reactive DV Abs on SARS-CoV-2 spike mutation(s), structural studies and docking simulations were performed using DV-2 envelope (E) Abs.

Signature mutation (s) in spike variants implied that majority of the above changes in the spike were driven by DV Abs rather than immune pressure of SARS-CoV-2 (preceding variants). This was supported by the fact that the lately emergent SARS-CoV-2 variants like Omicrons (2022-23) were at least 50% less cross-reactive to DV compared to preceding predominant strains.

This was further supported by the observed cross-binding of pre-pandemic DV Ab-positive serums to spike synthetic peptides in ELISA, designed from certain regions of the spike RBD which showed maximum cross-interactions with DV E Abs.

## 1. Introduction

From the beginning of the COVID-19 pandemic, a unique epidemiological observation became increasingly relevant and prominent in the midst of a significant public health emergency. Disease severity and mortality were observed to be significantly lower in DV endemic regions compared to DV non-endemic parts during COVID-19[1]. Subsequent research revealed that Abs against DV, a member of *Flaviviridae* family of viruses, had the ability to cross-react with SARS-CoV-2 and *vice versa*[2][3][4]. We furnished multiple pieces of evidence that DV Abs could also “cross-neutralize” betacoronaviruses under *Coronaviridae* family in cell culture, including SARS-CoV-2 Wuhan-hu-1 (WT)[5].

The inverse association between these two viruses from two separate families has now been reported from several countries[3][6][7]. Our previous computational investigations rightly indicated that DV-2 E Abs had high potential to bind to the Receptor-Binding Domain (RBD) of the SARS-CoV-2 wild-type (wt) spike (S) protein, which included several S protein residues involved in the host ACE2 receptor binding[8].

Conversely, *in vitro* investigation by others have confirmed that SARS-CoV-2 Abs raised against spike subunit 1 (S1)-RBD cross-reacted with DV E protein and reduced dengue pathogenicity in animal model[9]. Our laboratory had already shown that the DV cross-reacting SARS-CoV-2 Abs could effectively neutralise DV-1-4 in cell culture based virus neutralization assays (VNTs)[4][9][10]. This “cross-neutralization” potential suggests that the two viruses may be able to cross-protect one another. Observations during the active years of pandemic (2020-21) indicated that COVID-19 was less severe in DV endemic areas and *vice versa*, probably due to mutual cross-protection. In our ongoing efforts to monitor the dynamics of this “dengue and COVID-19 conundrum,” we had summarised many other evidence of mutual cross-interaction and their implications on public health in our previous review [11].

However, towards the end of 2020 when the first wave was waning[10] (see Fig. 3[10]), the situation in DV endemic areas changed when new mutations in the SARS-CoV-2 genome began to emerge[12]. Because of their lack of proof-reading function, RNA viruses are more susceptible to mutations[13]. Although SARS-CoV-2 has some proof-reading machinery, numerous mutations began to progressively accumulate in its genome (Supplementary Table 1). Spike protein is one of the most abundant proteins in SARS-CoV-2; structural in nature and a significant proportion of serum Abs are directed against spike [14]. New spike variants emerged over time; accumulating increasing number of mutations including insertions and deletions (the Omicrons) and breaching the prior variants’ serological protectivity[15]. The structural alteration of spike protein appeared to play an important role in this.

These new mutations in the emerging variants manifested a unique immunological escape potential besides other properties[16], which often contributed to serious health crisis in places that were previously unaffected by the first wave of COVID-19, i.e. DV endemic regions like India and Brazil. India, as a prime DV affected region, experienced a multi-dimensional health scarcity during the Delta wave of COVID-19 from April to June 2021[17]. So, the earlier DV-mediated cross-protectiveness against SARS-CoV-2 infection raised several questions: Are the variants getting selected to DV immune pressure besides preceding SARS-CoV-2 variant(s) immunity; if so, is the potential of DV immunity to cross-protect against mutated spike variants getting lessened due to the immune escape? Are selected escape mutations imparting more virulence and/or transmissibility to the emerging variants? We know that Abs exert selection pressure on the wt pathogen to prevent subsequent infection. Due to quasi-species nature of RNA viruses, spontaneous mutations of target antigen (s) of the pathogen get selected against existing immune pressure, allowing SARS-CoV-2 to reduce/overcome the protective barrier[18]. Besides immune escape, these mutations can also positively or negatively impact on virus fitness and pathogenicity.

In the present study, we examined how earlier highly cross-reactive DV Abs [4] might have positively influenced the spike protein evolution in DV endemic areas towards acquiring more mutations. There exists at least one other previously published meta-analysis report that suggested that patients co-infected with SARS-CoV-2 and DV, showed more amino acid substitutions in SARS-CoV-2 compared to DV negative group[19].

We have also surveyed the literature on mutations of SARS-CoV-2 spike protein, the major antigen of this virus, concentrating on their impacts on antigenicity and contextualising them in the protein structure. We created structural profiles of 5 mutant variant spike protein by incorporating in the Wuhan-hu-1 (WT) spike structure, all accessible and reported signature mutations characteristic for major variants namely B.1.1.7 (Kent), B.1.617 (Delta), B.1.617.2.1 (Delta-plus), and B.1.1.529 (Omicron-BA.1 and BA.2 variants). We also docked the mutated spike variants with DV-2 E Abs using the FFT-based global docking servers namely the ClusPro and HDOCK. Our computational studies and ELISA-based S peptide and DV-Abs binding assays provided robust evidence that the cross-reactive DV Abs have posed selection pressure on SARS-CoV-2 spike resulting in emergence of SARS-CoV-2 variants with increasing number of amino acid mutations and deletions in the exposed S1 protein and these emerged late variants like Omicrons became indeed, significantly less cross-reactive to DV[10].

## 2. Results

### 2.1. Structural change in the receptor-binding domain was observed in the mutated variants of the SARS-CoV-2 spike protein monomer

The I-TASSER-predicted[20][21] structures of the three SARS-CoV-2 variants viz. B.1.1.7 (Kent), B.1.617 (Delta) and B.1.617.2.1 (Delta-plus) showed the RBD in “lying down” conformation (Fig. 1a). In this connection, it was important to note that the ectodomain of SARS-CoV-2 wt spike protein is a class 1 fusion protein (approximately 1200 aa long) which constantly changed its conformational rearrangement position from RBD ‘up’ to ‘down’ and sometimes *vice versa* [22]. Previously, the X-ray crystallographic data of wt spike protein (PDB ID: 6VSB) presented RBD in “up” conformation. The “up” conformation was readily accessible for neutralization with monoclonal Abs and for tight and fast binding with ACE2 [23]. Here I-TASSER predicted models for four of the variants, namely, B.1.1.7, B.1.617, B.1.617.2.1 and B.1.1.529 (Omicron-BA.2) consistently depicted inherent RBD in “down” conformation with increased RMSD value (Å) (Root mean square deviation) when aligned with Wuhan-hu-1(WT) spike protein (Fig. 1b).

**Fig. 1.**
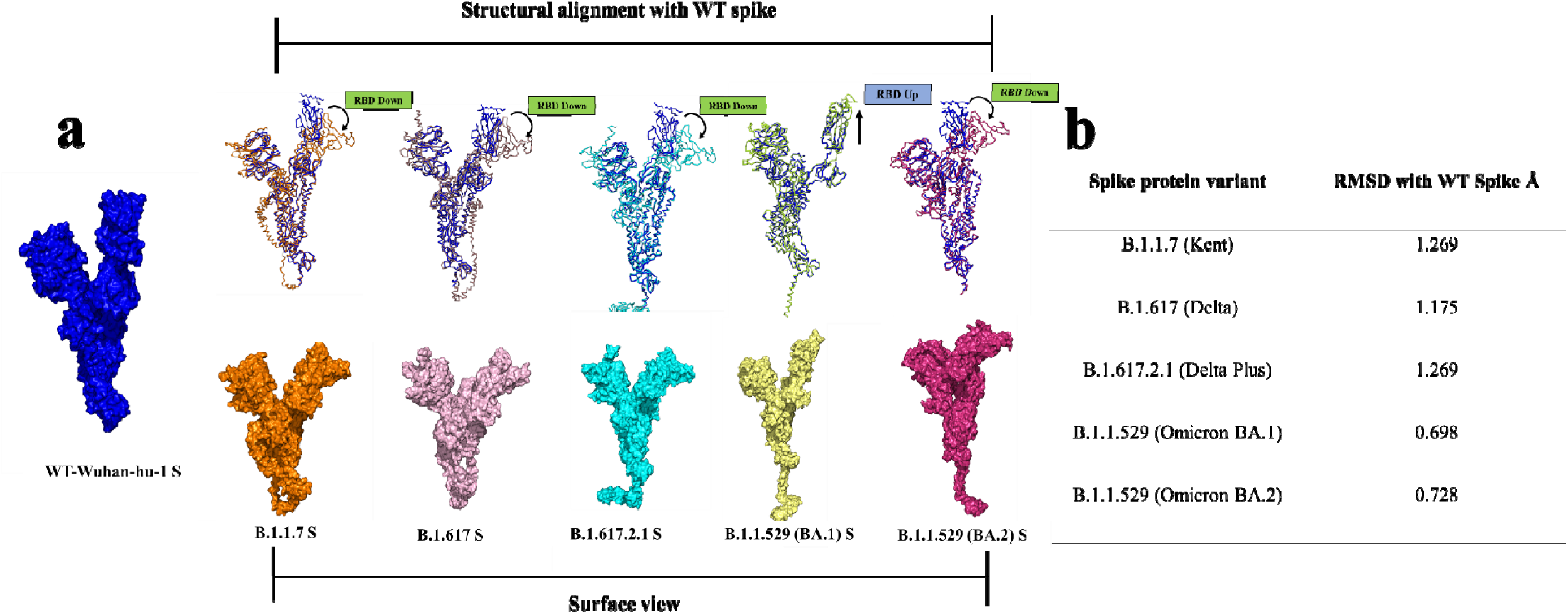
RBD conformation paradigm of mutated spike proteins and RMSD calculation with Wuhan-hu-1 (WT) spike protein. **a.** Surface view of each I-TASSER predicted mutant variant spike protein. Structural alignment of mutant spikes: B.1.1.7 (Kent), B.1.617 (Delta), B.1.617.2 (Delta-plus) and B.1.1.529 (Omicron-BA.2) with Wuhan-hu-1(WT) spike showed RBD in “down” conformation. In case of B.1.1.529 (Omicron BA.1), the RBD was found to be in “up” conformation. **b.** RMSD values (Å) of each mutated spike protein after structural alignment with Wuhan-hu-1(WT) spike protein.

The structural deviations were observed mainly in the RBD region of the mutated proteins. The predicted data correlated with the experimental evidence regarding conformational paradigm of spike protein[24][25][26]. This inherent structural shift is comprehendible as these mutational changes might have contributed to the “lying down’ position of spike protein RBD allowing for evasion of immune response to wt spike or vaccine Abs[27]. Thus, decreased effectiveness of COVID-19 vaccines against the variants[28] could be consequential to the mutation(s)-driven structural changes.

On the other hand, the I-TASSER-predicted spike structure of B.1.1.529 (Omicron-BA.1) showed the RBD “up” conformation and low RMSD after structural alignment with Wuhan-hu-1(WT) spike. This finding also correlated with experimental findings[29] where electron microscopy data on Omicron-BA.1 RBD structure supported that it was in “up” conformation.

### 2.2. *In silico* docking of dengue virus antibodies with SARS-CoV-2 spike protein mutants produced decreased RBD interaction compared to wild-type SARS-CoV-2 spike protein

For antigen-antibody docking analysis, available structure files of DV-2 E Abs [30] i.e., EDE1 C8, EDE2 A11, EDE2 B7 and EDE1 C10 were docked with the I-TASSER predicted structure files of the spike protein of the variants (B.1.1.7, B.1.617, B.1.617.2.1, B.1.1.529-BA.1 and BA.2) in two FFT-based global docking servers HDOCK and ClusPro. Among all predicted structures of each mutant spike, only those with high C-score, were used in further docking analysis. Following the docking exercise, 10 structures were retrieved from each server for each antigen-antibody docking pair. Total 80 predicted docked structures were analyzed for each of the 5 mutated spike variant(s) docked with each of the 4 DV-2 E Abs. First, the docking frequency (DF) in RBD of spike was calculated for each antigen-antibody pair (Fig. 2a). Thereafter, cumulative docking frequency (CDF) was calculated adding the DF of all 4 DV-2 E Abs for one particular spike variant (Fig. 2b). The amino acid boundaries of the RBD region of the Wuhab-hu-1 (WT) spike protein were used as a reference to draw the RBD regions of all spike mutants throughout the study. In comparison to the previously published docking experiment with Wuhan-hu-1 (WT) spike[8], it was found that CDF of DV-2 E Abs with each successively emerged SARS-CoV-2 mutant was progressively less, in general.

**Fig. 2.**
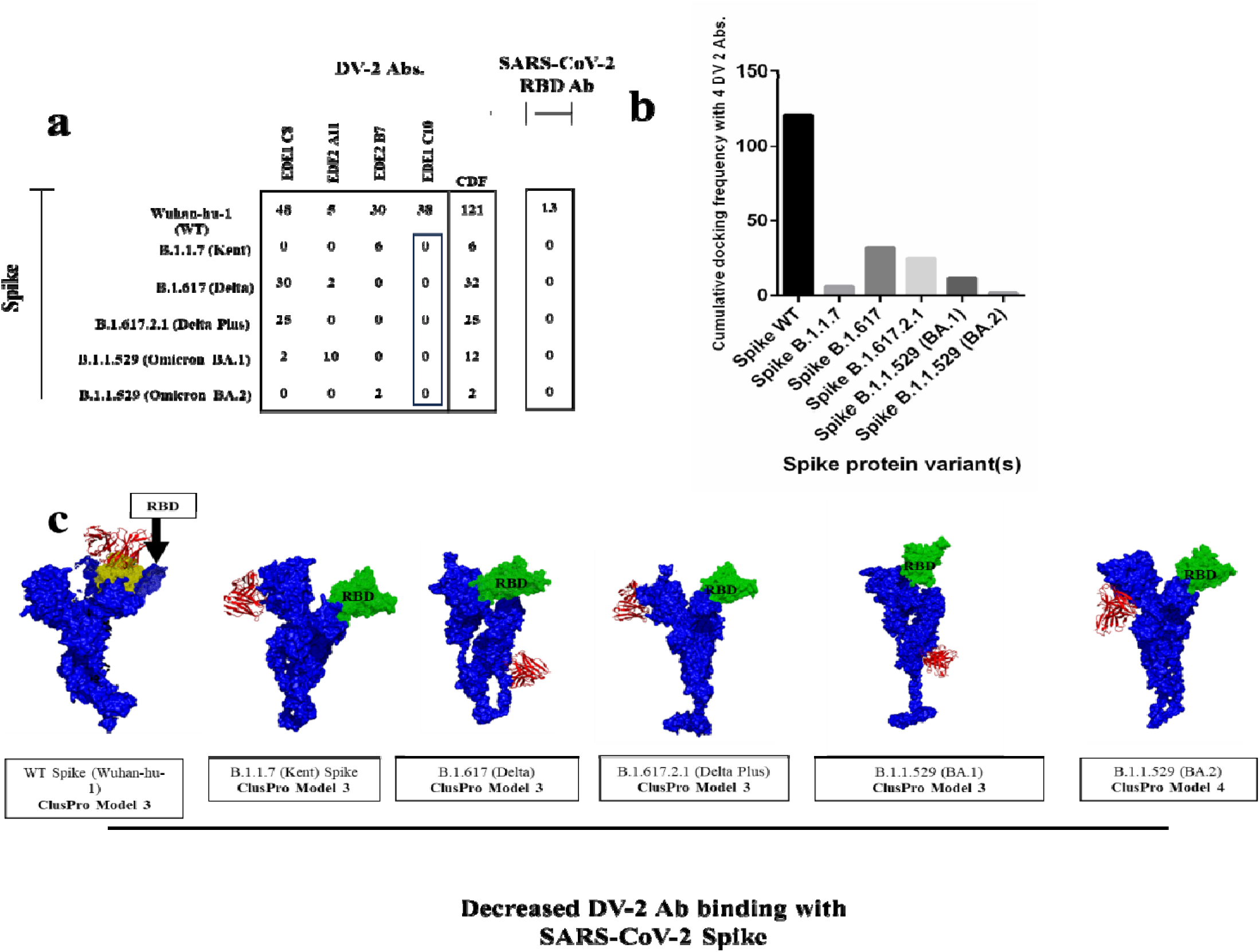
*In silico* docking of DV-2 E Abs with predicted SARS-CoV-2 mutant variant(s) **a.** Docking frequency (179 DF) table of the binding of 4 DV-2 E Abs and SARS-CoV-2 RBD Ab in the RBD of SARS-CoV-2 wild-type (Wuhan-hu-1), B.1.1.7 (Kent), B.1.617 (Delta), B.1617.2.1 (Delta-plus) and B.1.1.529 (Omicron-BA.1 and BA.2) spike. **b.** 2D bar graph of cumulative docking frequency (CDF) depicting lower interaction frequency of DV-2 E Abs with SARS-CoV-2 mutant spikes. **c.** When docked with the wild-type Wuhan-Hu-1 spike protein, representative docking models of one DV-2 E Ab, EDE1 C10, were found to have neutralising activity (RBD compromised with Ab; highlighted yellow). However, the same antibody did not attach to the RBD area of any of the mutant spikes (RBD free; highlighted green).

Previously, the DV-2 E Abs tended to interact in the RBD (receptor binding domain; aa 333-527) of the wt spike protein with a CDF of 121 involving all the 4 DV-2 E Abs (48 DF with EDE1 C8 + 5 DF with EDE2 A11 + 30 DF with EDE2 B7 + 38 DF with EDE1 C10)[8]. For the mutated variants, however, it was found that there was only DF of 6 between the EDE2 B7 and the B.1.1.7 (Kent) spike, while the remaining 3 Abs showed no DF in the RBD region. The B.1.617 (Delta) spike showed a small amount of interaction, with a DF of 30 in the RBD area with EDE1 C8 and DF of 2 with EDE2 A11. This resulted in a CDF of 32 for B.1.617 with DV-2 E Abs. For the B.1.617.2.1 (Delta plus) spike, a DF of 25 was observed with only EDE1 C8 in RBD. In the case of the B.1.1.529 (Omicron-BA.1), a DF of 2 was found with EDE1 C8, and DF of 10 was found with EDE2 A11 resulting in a CDF of 12. Similarly, in case of B.1.1.529 (Omicron-BA.2) a DF of only 2 was found with EDE2 B7. It was noteworthy that the successively emerged variants that evolved in DV endemic regions (Deltas and Omicrons), showed progressively reduced binding to DV-2 E Abs providing evidence of positive selection under DV Abs pressure with gradual accumulation of mutations in the spike protein, besides insertions and deletions (Supplementary Table 1). The Kent variant originated in UK (DV non-endemic) from the Wuhan-hu-1 (WT) strain around September 2020 and showed a low CDF of 6.

Out of the 4 DV-2 E Abs, only EDE1 C10, did not interact with any of the mutant variants (Fig. 2c). Representative docking image clarified how the RBD of wt spike protein was previously compromised (marked yellow) by the same antibody EDE1 C10[8]. Gradually, the mutations in the wt spike and the resultant structural shift in the RBD had taken toll on the cross-reactivity potential of the DV-2 E Abs with the emergent SARS-CoV-2 variants. For EDE1 C10 docking with the mutated spike of SARS-CoV-2 variants, no RBD interaction (marked green) was observed, leading to full exposure of the RBD for interaction with ACE2. This could be one reason for high transmissibility of the SARS-CoV-2 emergent variants like the Omicrons[31][32]. Detailed list of docked amino acids in the RBD region of each spike protein variant for each DV-2 E Ab can be found in the Supplementary excel information (S1-5, Mendeley Data: https://data.mendeley.com/datasets/gvvb4rf98h/1).

Structure files of all the docked complexes can be found in the online repository site [Mendeley Data: https://data.mendeley.com/datasets/gvvb4rf98h/1]. As a positive control approach to our docking study, we had also done the docking simulations between SARS-CoV-2 Wuhan-hu-1 (WT) RBD-directed Ab (PDB ID: 7BWJ) and SARS-CoV-2 spike mutants and Wuhan-hu-1 (WT) spike [Fig. 2a, Supplementary excel information S6, Mendeley Data]. Except Wuhan-hu-1 spike, the RBD directed Ab didn’t dock with any of the predicted mutant spike protein structures supporting the emergent variants as Ab escape mutants. Findings of this study corroborated with experimental observations that confirmed that Delta and Omicron were indeed, escape variants to Abs elicited by preceding variants and even vaccines, [33][34][16][35] further strengthening the reliability and robustness of our predictions.

### 2.3. Trend of DV-2 E Abs pressure on SARS-CoV-2 spike RBD mutation (s)

The marked spike amino acid sequence of each mutated SARS-CoV-2 variant provides an insight of the trend of DV-2 E Abs pressure on the selection of amino acid mutations in the variants within the RBD and RBM (Receptor-Binding Motif) regions of spike S1 (Fig. 3a and 4). The yellow marked region in the top number scale of spike amino acid sequence (Fig.3a and 4) represents the RBD region and the light blue colored region within RBD, marks the RBM region of each variant. The amino acid boundaries of the RBD and RBM have been marked, using the amino acid boundaries of the Wuhan-hu-1 (WT) spike RBD and RBM as a reference [8].

**Fig. 3.**
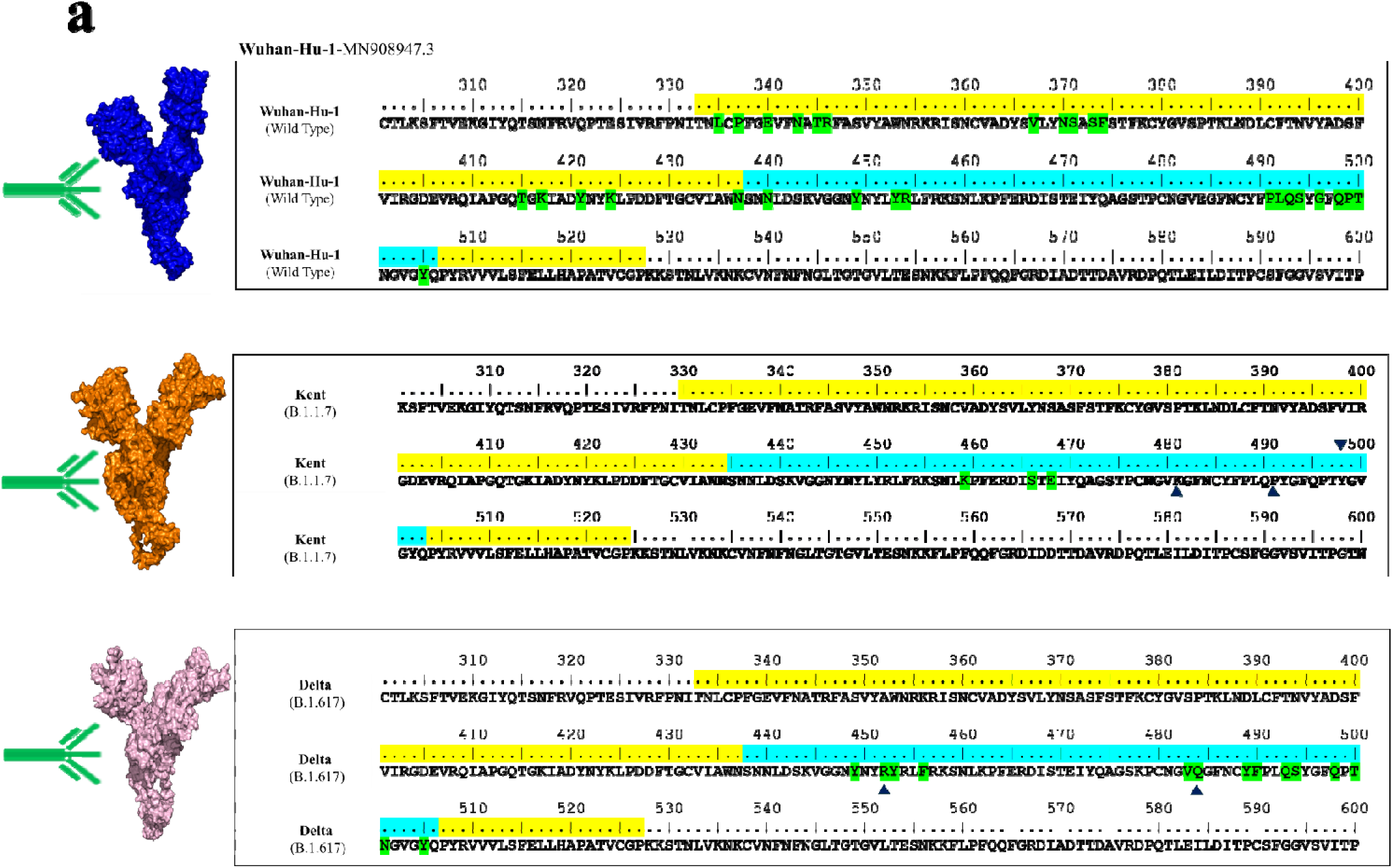
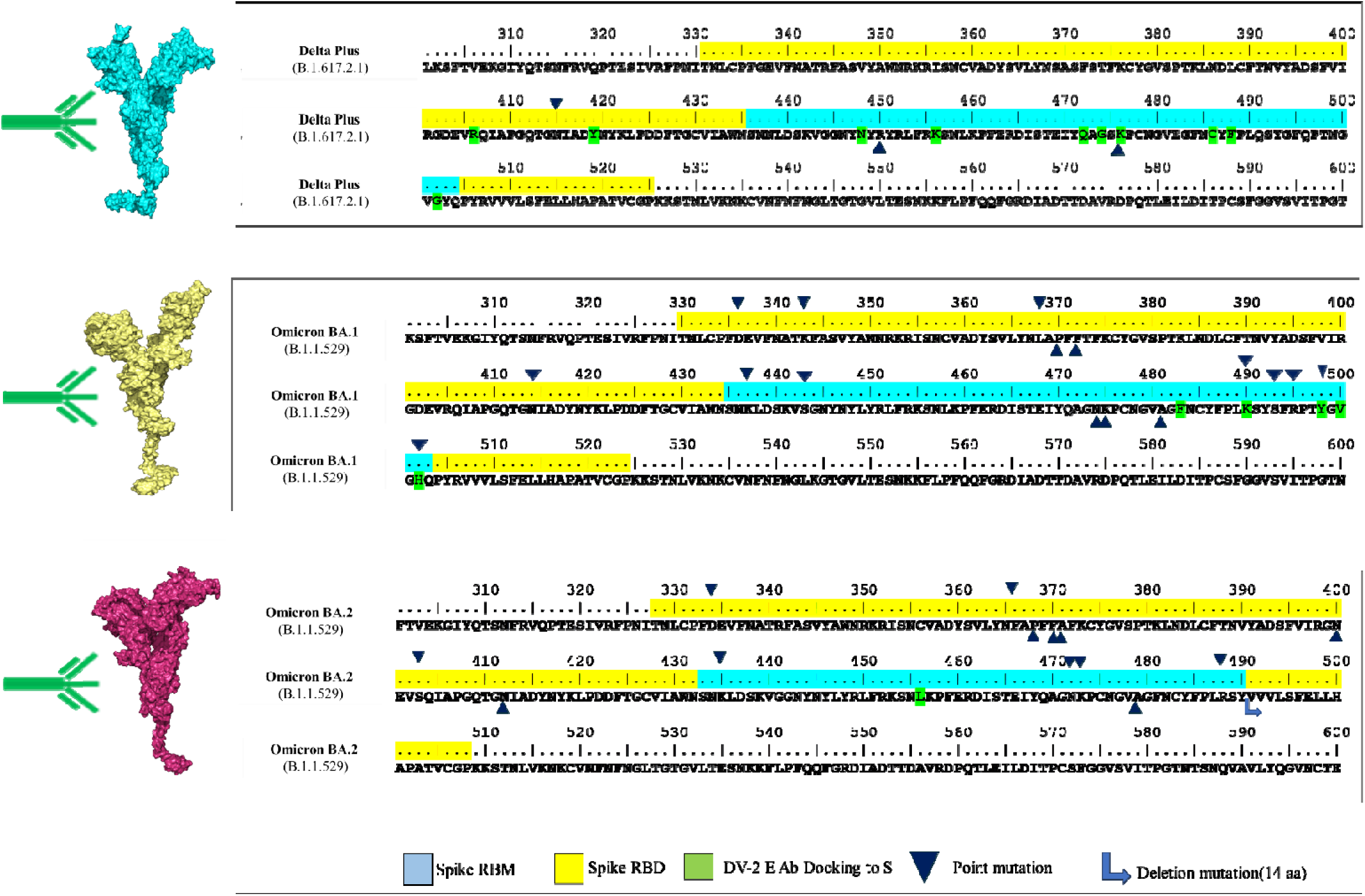
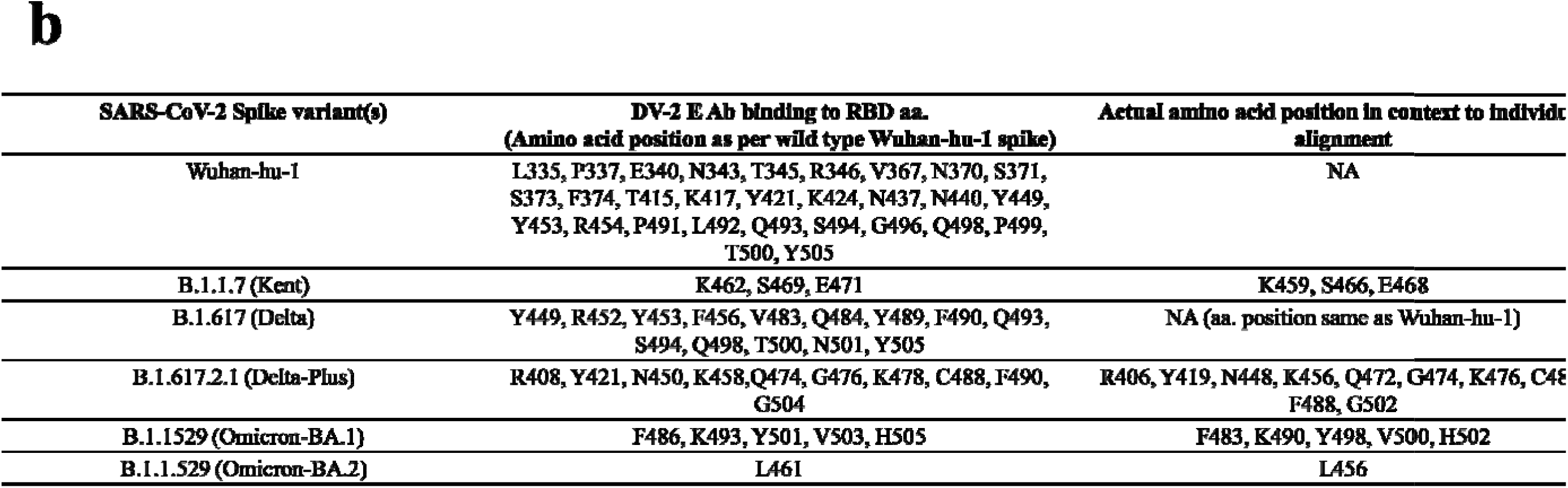
RBD amino acid sequence of SARS-CoV-2 spike variants showing reduced interaction with DV-2 E Abs compared to wild type spike RBD. **a.** Predicted interaction sites of DV-2 E Abs with SARS-CoV-2 spike variants. Yellow marked region denotes RBD region of each mutant spike. Light blue marked region denotes RBM region of each mutant spike. Amino acids marked in green are those which docked with DV-2 E Abs. Dark blue arrow and light blue arrow and rectangle indicate point mutation and deletion mutations respectively **b.** List of amino acids of each mutant spike which interacted with DV-2 E Ab.

**Fig. 4.**
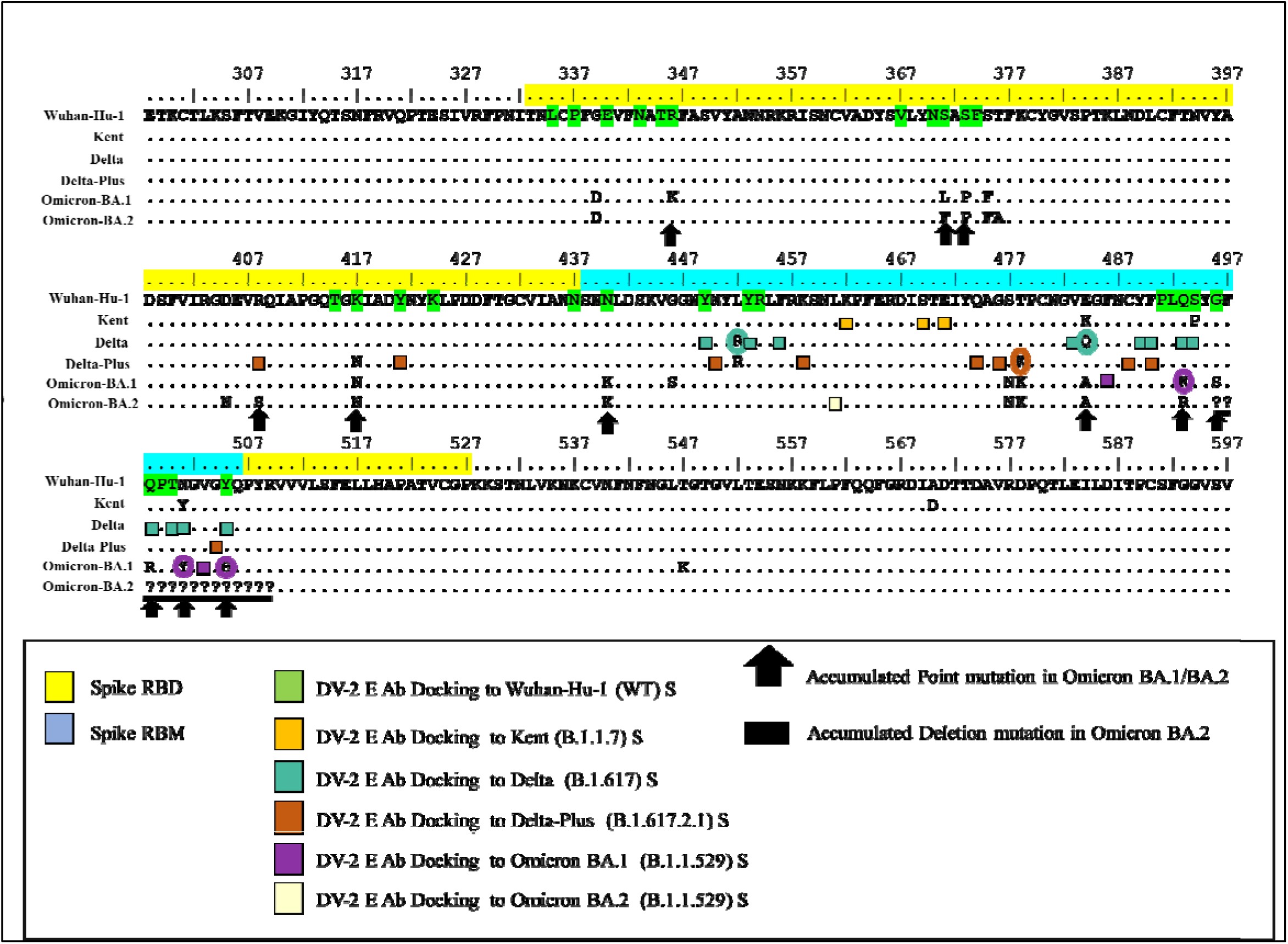
Multiple sequence alignment view showing how DV E Abs might have imparted selection pressure in spike protein RBD region contributing to emergence of SARS-CoV-2 variants with increasing number of mutations/deletions. From SARS-CoV-2 Wuhan-hu-1(WT) spike to SARS-CoV-2 B.1.1.529 (Omicron-BA.2) spike DV E Abs interaction appeared to play an instrumental role in selecting mutation (s) in RBD region. In the alignment, yellow coloured bar over the alignment signifies the border of RBD region. The light blue coloured rectangle denotes the RBM region. Different coloured boxes indicate different variant(s) spike amino acid residues predicted to dock/bind with DV-2 E Abs. The black arrows signify point mutations selected and accumulated in B.1.1529 (Omicron-BA.2) under DV Ab pressure. The black rectangle denotes stretch of 14 amino acids deleted in Omicron-BA.2.

Green marked amino acids are the ones that showed interaction with the DV-2 E Abs (Fig. 3a). It is clearly visible from this docking study that both the amino acid numbers and their CDF with DV-2 E Abs in RBD region of the spike mutants have markedly reduced over time in case of successively emerged variants.

Previously, 29 amino acids docked with DV-2 E Abs, but this number has steadily decreased in subsequent versions. It reached its lowest point in Omicron-BA.2, where only one amino acid interacted with DV E Abs in the said regions (Fig. 3b).

Cluster of colored boxes depicts interaction site of DV-2 E Abs in the multiple sequence alignment picture (Fig. 4). It portrays a visual representation of DV E Abs’ pressure on SARS-CoV-2 spike protein RBD. For instance, 14 RBD amino acids were identified where DV-2 E Abs had interacted *in silico* with Wuhan-hu-1(WT) or any other successive variants up to Delta-plus, which have resulted in amino acid mutation or deletion(s) in the lately emergent B.1.1.529 (Omicron-BA.1 and/or BA.2) (Table 1). Out of these 14 amino acids, 6 were ACE2 interaction sites [8]. So, it is imperative from our computational studies that DV-2 E Abs might be exerting a pressure for selection of crucial mutations and emergence of SARS-CoV-2 variants in dengue endemic regions. Also, it is apparent from the multiple sequence alignment map (Fig 4) of B.1.1.529 (Omicron-BA.2) that after experiencing repeated docking of DV-2 E Abs in the region spanning 496-509 (Fig. 4) aa in previous variants, a 14 amino acid deletion emerged. This deleted region exhibited 4 DV Abs-interacting amino acids for Wuhan-hu-1 (green), Delta (deep blue) and Delta-Plus (brown) and 5 mutations in case of BA.1. Three of these mutations (namely Q498R, N501Y and Y505H) occurred from amino acids that interacted with DV Abs in preceding variants like Wuhan-hu-1, Delta and Delta-Plus. Among the 21 most common amino acid mutations found in the RBD region of both Omicron-BA.1 and BA.2 spike, 13 (61.9%) occurred at positions where DV-2 E Abs had been predicted to bind to the spike protein in earlier variants.

**Table 1.** List of mutated RBD amino acids in B.1.1.529 (Omicron-BA.1 and BA.2) in DV-2 E Ab docking site. The left column shows a list of RBD amino acids that interacted with DV-2 E Abs when docked with Wuhan-hu-1 or another spike variant. Amino acids demarcated in bold letter denote ACE2-interacting residues [7]. The next column shows a list of subsequent mutated RBD amino acids in B.1.1.529 (Omicron-BA.1 and BA.2). The next column denotes the genomic codon mutation of subsequent amino acid mutations. The subsequent column denotes the type of genomic mutations observed. The last column denotes the impact on DV-2 E Abs docking due to mutation.

| Name and position of amino acids in WT spike or any variant spike which interacted with DV-2 E Abs in silico | In silico interaction sites show mutations in Omicron BA.1 and BA.2 | Genomic mutation | Mutation type | Effect in DV-2 E Abs docking |
| --- | --- | --- | --- | --- |
| R346 [Wuhan-hu-1] | K346 [Omicron BA.1] | AGA→AAA [Omicron BA.1] | Transition [Omicron BA.1] | Non-reactive |
| S371 [Wuhan-hu-1] | L371 [Omicron BA.1]<br>F 371 [Omicron BA.2] | TCC→ CTC [Omicron BA.1]<br>TCC→ TTC [Omicron BA.2] | Transition [Omicron BA.1]<br>Transition [Omicron BA.2] | Non-reactive |
| S373 [Wuhan-hu-1] | P373 [Omicron BA.1]<br>P 373 [Omicron BA.2] | TCA → CCA [Omicron BA.1]<br>TCA→ CCA [Omicron BA.2] | Transition [Omicron BA.1]<br>Transition [Omicron BA.2] | Non-reactive |
| R408 [Delta Plus/ B.1.617.2.1] | S408 [Omicron BA.2] | AGA → AGC [Omicron BA.2] | Transversion [Omicron BA.2] | Non-reactive |
| K417 (ACE2 interacting aa.) [Wuhan-hu-1] | N417 [Omicron BA.1]<br>N417 [Omicron BA.2] | AAG→AAT [Omicron BA.1]<br>AAG→AAT [Omicron BA.2] | Transversion [Omicron BA.1]<br>Transversion [Omicron BA.2] | Non-reactive |
| N440 [Wuhan-hu-1] | K440 [Omicron BA.1]<br>K440 [Omicron BA.2] | AAT→AAG [Omicron BA.1]<br>AAT→AAG [Omicron BA.2] | Transversion [Omicron BA.1]<br>Transversion [Omicron BA.2] | Non-reactive |
| Q484 [Delta/ B.1.617] | A484 [Omicron BA.1]<br>A484 [Omicron BA.2] | CAA→GCA [Omicron BA.1]<br>CAA→GCA [Omicron BA.2] | Transversion [Omicron BA.1]<br>Transversion [Omicron BA.2] | Non-reactive |
| Q493 [Wuhan-hu-1, Delta/B.1.617]<br>K493 [Omicron BA.1] | K493 [Omicron BA.1]*<br>R493 [Omicron BA.2] | *Sequence could not be retrieved from database, but mutation was reported in Omicron BA.1*<br>CAA→CGA [Omicron BA.2] | NA<br>Transition [Omicron BA.2] | Reactive in Omicron BA.1<br>Non-reactive in Omicron BA.2 |
| G496 (ACE2 interacting aa.) [Wuhan-hu-1] | S496 [Omicron BA.1]<br>Deleted [Omicron BA.2] | GGT→AGT [Omicron BA.1]<br>Deleted [Omicron BA.2] | Transition [Omicron BA.1]<br>Deletion [Omicron BA.2] | Non-reactive |
| Q498 (ACE2 interacting aa.) [Wuhan-hu-1, Delta/B.1.617] | R498 [Omicron BA.1]<br>Deleted [Omicron BA.2] | CAA→CGA [Omicron BA.1]<br>Deleted [Omicron BA.2] | Transition [Omicron BA.1]<br>Deletion [Omicron BA.2] | Non-reactive |
| T500 (ACE2 interacting aa.) [Wuhan-hu-1, Delta/B.1.617] | Deleted [Omicron BA.2] | Deleted [Omicron BA.2] | Deletion [Omicron BA.2] | Non-reactive |
| N501 (ACE2 interacting aa.) [Delta/B.1.617] | Y501 [Omicron BA.1]<br>Deleted [Omicron BA.2] | AAT→TAT [Omicron BA.1]<br>Deleted [Omicron BA.2] | Transition [Omicron BA.1]<br>Deletion [Omicron BA.2] | Reactive in Omicron BA.1<br>NA [Deleted in Omicron BA.2] |
| V503 [Omicron BA.1] | Deleted [Omicron BA.2] | Deleted [Omicron BA.2] | Deletion | NA [Deleted in Omicron BA.2] |
| Y505 (ACE2 interacting aa.) [Wuhan-hu-1, Delta/B.1.617] | H505 [Omicron BA.1]<br>Deleted [Omicron BA.2] | TAC→CAC [Omicron BA.1]<br>Deleted [Omicron BA.2] | Transition [Omicron BA.1]<br>Deletion [Omicron BA.2] | Reactive in Omicron BA.1<br>NA [Deleted in Omicron BA.2] |

\*Q493K mutation in Omicron-BA.1 was reported in many articles[36], but the sequence was not found from any gene repository database where this mutation persists.

Of the 17 amino acid substitutions in Omicron-BA.1 and BA.2 that showed DV-2 Ab docking to the wt or any preceding variant amino acid (i.e. suggestive of DV Ab selection pressure), 10 mutations were due to single base transition and remaining 7 were because of single base transversion (Table 1).

### 2.4. SARS-CoV-2 peptides cross-reacted with dengue pre-pandemic serum samples

Depending on increased frequency of predicted DV-2 E Abs-interactions with the spike residues, four stretches of spike amino acids were chosen and synthesized to test their actual cross-reactive/binding potential with DV Abs. One such peptide, CoV-2 Pep-1 (NLCPFGEVFNATRFA) contains a previously reported cross-reactive region FNATR [9]. This epitope was identified by DV cross-reacting SARS-CoV-2 RBD IgG from a 12-mer phage display library. This epitope was responsible for completely neutralizing cross-reactive SARS-CoV-2 Abs interaction with S1-RBD and DV E protein. Alongside this peptide, three more peptides were chosen which showed high degree of interaction with DV-2 E Abs as per our computational predictions of previous[8] and current study (Fig. 5a).

**Fig. 5.**
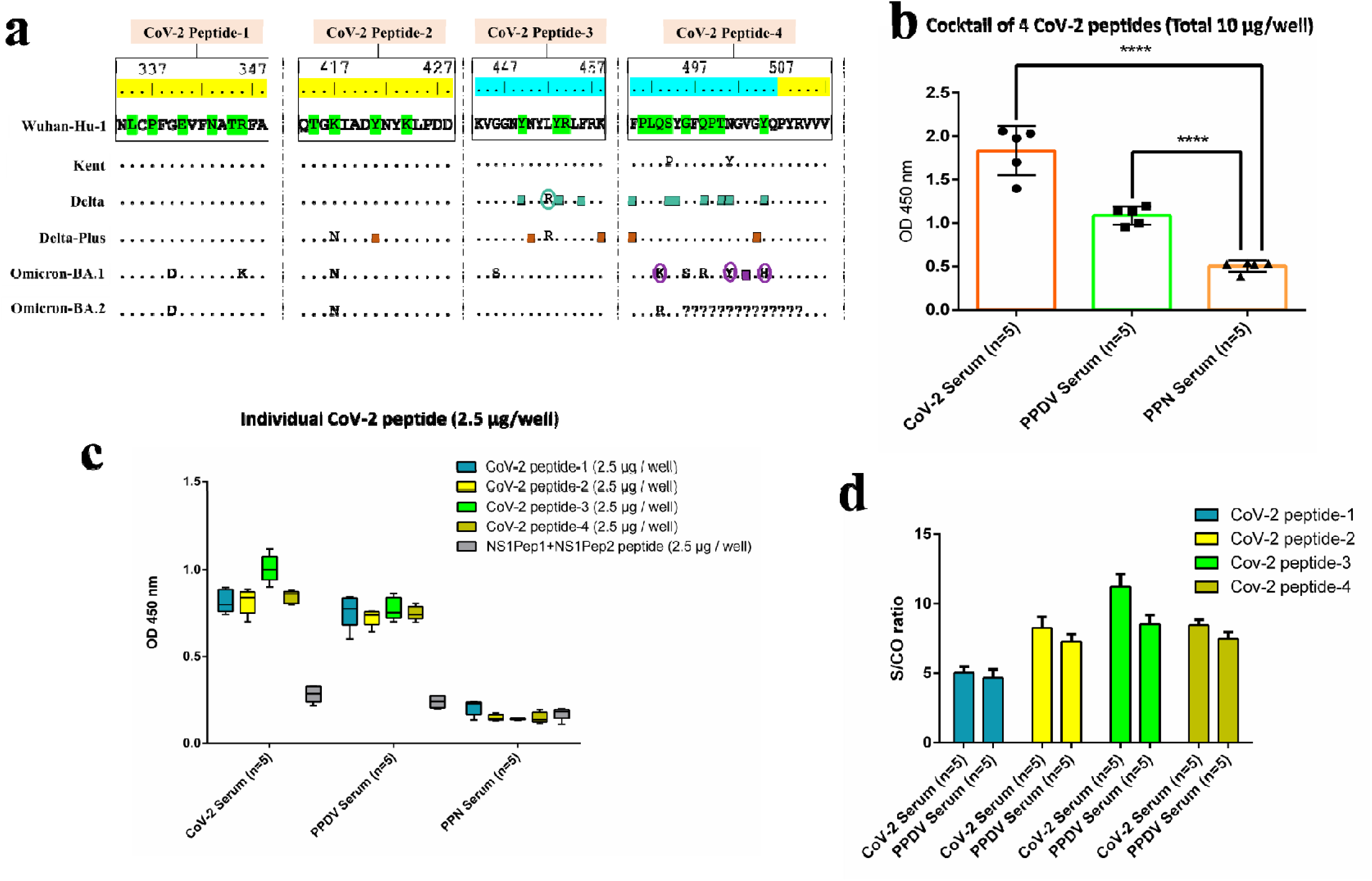
Truncated CoV-2 RBD peptide coating ELISA. **a.** Sequence of four CoV-2 peptides and their subsequent variant (s) sequence alignment. DV E Ab docking sites were marked as in Fig. 4. **b.** 2D bar graph of reactivity of all four CoV-2 peptides with CoV-2, PPDV, PPN serum samples (n=5/set). CoV-2 and PPDV serum samples had significantly higher reactivity (p < 0.0001) than PPN serum samples (negative control). p_Values were calculated from minimum five counts for each condition, by two tailed, unpaired t_test at 95% level of confidence. **c.** 2D bar graph of individual CoV-2 peptide 1-4 and NS1Pep1+NS1Pep2 peptide mix reactivity with CoV-2, PPDV and PPN serum samples (n=5/set); here CoV-2 peptide 1-4 showed higher reactivity with COV-2 and PPDV serums compared to PPN serum samples. NS1Pep1+NS1Pep2 peptide mix had significantly lower reactivity with all serum samples. **d.** 2D bar graph showing the S/CO values of individual CoV-2 peptide reactivity with CoV-2 and PPDV serum samples (n=5/set). Each column represents mean ± SD from five replicates for each test serum sample.

At first, all the four peptides were coated per well of 96 well ELISA plate (2.5 µg per peptide, total 10 µg/well). Five archived 2020-21 pandemic COVID-19 serum samples (CoV-2 serum samples) reacted with the peptides mix, thereby acting as effective positive control of the assay. Five pre-COVID-19 pandemic DV infected serum samples (PPDV serum samples) also cross-reacted with these peptide stretches. Their serological status revealed that they contained DV IgG/IgM and faintly cross-reacted in SARS-CoV-2 IgG/IgM LFIA tests (Fig 5b). Pre-COVID-19 pandemic normal serum samples (PPN serum samples) which had no detectable SARS-CoV-2 or DV IgG/IgM, didn’t react significantly with these four-peptides mix. They served as the negative control of the study (Fig. 5b).

In subsequent ELISA assays, when these four peptides were coated individually in each well of 96 well plate (2.5 µg/well), it was observed that all of these peptide stretches individually also reacted significantly with both CoV-2 and PPDV serum samples compared to PPN serum samples (Fig. 5c). As negative control of the peptide reactivity, we also used mix of two DV1 NS1 truncated peptide stretches (NS1Pep1+NS1Pep2).

A two-way ANOVA revealed significant effects of serum type and peptide type on OD responses in the ELISA assays. The analysis revealed significant main effects of both serum type (F (2, 60) = 878.81, p < 0.0001) and peptide type (F (4, 60) = 144.34, p < 0.0001), along with a significant interaction effect (F (8, 60) = 41.95, p < 0.0001).

Multiple t-tests with Bonferroni correction further demonstrated that CoV-2 serum samples showed significantly higher reactivity compared to PPDV serum samples for peptides 2, 3, and 4 (p < 0.002), and markedly higher reactivity than the PPN serum samples for all four peptides (p < 0.0001). Additionally, PPDV serum samples exhibited significantly higher OD responses than PPN serum samples for peptides 1–4 (p < 0.0005), supporting cross-reactivity.

Peptide 3 displayed the highest reactivity overall (t = 6.71, p < 0.0001), highlighting its immune-dominant nature (5c, d). No significant differences were observed in the reactivity of NS1Pep1+NS1Pep2 peptides across CoV-2, PPDV, and PPN serum samples (p > 0.05), confirming their role as negative controls. These results indicated that the CoV-2 test serum samples contained SARS-CoV-2 Abs against all these four peptides and the PPDV serum samples, indeed cross-reacted with these antigenic COV-2 spike peptides. The S/CO values of Abs-binding to individual peptides also gave an indication of the comparative reactivity of CoV-2 and PPDV serum samples with the four peptide stretches. It was observed that CoV-2 peptide 3 had slightly higher reactivity with both the populations of serum samples with respect of other three peptide stretches (Fig. 5d).

## 3. Discussion

Cross-reactivity between DV and SARS-CoV-2 during the early waves of COVID-19 was clearly demonstrated to have an important role in reciprocal cross-protection and impact on disease progression in DV endemic regions globally. As mentioned before, in dengue endemic regions like the Indian sub-continent and Southeast Asia, COVID-19 severity and mortality were significantly lower compared to dengue non-endemic regions during the active years of the pandemic and *vice versa* (2020-21).

The COVID-19 death toll per million population in India, a DV endemic country, was at least 10-fold less compared to western non-DV endemic regions like North America and Europe[37]. Lately, we have published our research findings to consolidate the above observation. We had previously demonstrated that pre-pandemic DV Ab-positive serum samples were nearly 38% cross-reactive in SARS-CoV-2 IgG/IgM serological tests [2], supported by similar studies from Israel and Taiwan [3][9]. We further showed that DV Ab-positive human serum samples could indeed, “cross-neutralize” (hence have the potential to cross-protect) betacoronaviruses like MHV-1 in VNT in cell culture and SARS-CoV-2 (Wuhan-hu-1) in surrogate VNT (Ab mediated inhibition of RBD-ACE2 interaction) in ELISA-based assay[5]. Further, a meta_analysis on the burden of respiratory viral diseases in different states of India carried out using data from 1970-2020 (fifty years) revealed that India has higher rates of infections of respiratory syncytial virus, influenza, parainfluenza and a trend of higher rate of rhinovirus infections[38]. But infection due to HCoVs was hardly reported from India. Even the Middle East Respiratory Syndrome_CoV (MERS_CoV) which caused nearly 487 deaths worldwide was never detected in India. Same is true for SARS_CoV which had few sporadic occurrences reported from the Indian subcontinent, a highly dengue endemic region [2][10]. The cross-immunity mediated protection due to DV appears to be there all the time against HCoVs in dengue endemic regions, in general.

It is therefore, obvious that SARS-CoV-2 must be evolving under the pressure of its own Abs and DV Abs. Incidentally, almost 50-90% Indians are seropositive for DV depending on regional endemicity[39] and DV IgGs generally do not wane with time like SARS-CoV-2 Abs[40][41][42]. That is the reason why pre-existing DV immunity can lead to antibody-dependent enhancement (ADE) due to subsequent infection with a different DV serotype[43][44]. In other words, long lived and persisting DV Abs is the major reason behind more severity generally observed in secondary dengue compared to primary infections [45].

In contrast, Abs due to SARS-CoV-2 or vaccines have been reported to hardly last longer than a year[46], resulting in worldwide reports of reinfections despite previous virus exposure/vaccination and necessitating booster vaccinations[47]. So, it may be argued that DV immunity posed greater selection pressure on SARS-CoV-2 over its own immunity. Thus, variants became more rapidly selected in DV prone regions, with increasing number of mutations in the structural proteins resulting in variants like Omicrons.

COVID-19 is hardly occurring in present times (2024) and does not appear to naturally spread in DV endemic regions; perhaps DV immunity has led to “melting down” of SARS-CoV-2 genetic information in case of the previously prevalent Omicron variants like BA.2 (with >30 mutations and deletion in RBD), resulting in “error catastrophe” that eventually culminates in virus extinction, as possibly happened in case of other HCoVs in DV prevalent regions, as discussed earlier. Again, during current times, COVID-19 is still spreading, albeit slowly and sporadically in dengue non-endemic countries like North America as SARS-CoV-2 immunity is weak and suboptimal, and DV immunity is absent. Rare incidences of COVID-19 in DV endemic regions are currently occurring most likely due to entry of variants like XBB from non-endemic regions by air and sea routes of travel[48][49].

Thus, our computation-based prediction results of SARS-CoV-2 Ab docking with each variant spike protein was consistent with experimental and epidemiological observations. Over time, Abs generated against the original SARS-CoV-2 Wuhan-hu-1 strain failed to block infection caused by subsequently emerged variants like Kent, Delta or Omicrons. The correlation of *in silico* and *in vitro* data significantly increased the robustness and reliability of our docking predictions. Our *in silico* docking with DV-2 E Abs revealed progressively lower degrees of spike protein interactions in case of the successively emerged variants compared to the Wuhan-hu-1 (WT) spike protein, the least with Omicron variant, BA.2 (last in our representative series).

Four peptide stretches of the SARS-CoV-2 Wuhan-hu-1 spike protein were synthesized. These regions were predicted to be the most targeted by DV-2 E Abs. Each peptide sequence was aligned with their mutated counterparts (Kent to Omicron-BA.2) to see successive docking by DV-2 E Abs (Fig. 5a). CoV-2 peptide 1 was found to contain 6 amino acids from the Wuhan-Hu-1 sequence targeted by DV-2 E Abs. CoV-2 peptide 2 had 4 such interacting amino acids from the Wuhan-Hu-1 sequence and 1 amino acid from the Delta Plus variant (total=5 interacting residues). CoV-2 peptide 3 included 3 amino acids from Wuhan-Hu-1, 4 amino acids from the Delta variant, and 2 amino acids from the Delta Plus variant (total=9 interacting residues). Finally, CoV-2 peptide 4 contained 9 amino acids from Wuhan-Hu-1, 7 from the Delta variant, 2 from Delta Plus, and 4 from the Omicron-BA.1 variant (total=22 interacting residues).

It appears that the peptide 4 stretch contained the cumulative highest number of residues under DV E Abs pressure, over the successively emerging variants from Wuhan-Hu-1 wt strain. That may be one strong reason why this stretch was selecting maximum mutations in the successively emergent variants. On top this stretch eventually showed 14 amino acids deletion in the lately emergent Omicron-BA.2 variant, which further suggests strong and persistent immune pressure over this region.

The results of the docking study suggested a high possibility of DV-2 E Abs cross-reacting with specific SARS-CoV-2 spike protein regions, potentially imparting immunological pressure. To confirm this, the four synthesized peptides were coated onto wells of a 96-well plate, and a standardized in-house ELISA was performed with three sets of serum samples. When the 10 µg peptide cocktail of 4 peptides was tested with SARS-CoV-2 Delta wave serums (positive controls), the samples showed high reactivity as expected. Pre-COVID-19 normal human serum samples showed significantly lower reactivity with the peptide cocktail. In contrast, pre-pandemic DV serum samples, which was also used in earlier assays to neutralize MHV-1[5], showed comparable reactivity close to the positive control OD values (Fig. 5b). When the peptides were tested individually at 2.5 µg /well with the same serum sets, all four peptides reacted with SARS-CoV-2 Delta wave serums and also cross-reacted with pre-pandemic DV serums. Pre-COVID-19 normal human serum continued to show lower reactivity with all four peptides. To assess specificity, DV NS1 peptide stretches NS1Pep1 and NS1Pep2 were tested (Fig. 5c). None of the serum samples reacted with these NS1 peptides, confirming the assay’s specificity. Further analysis of the S/CO ratios from individual peptide ELISA tests identified CoV-2 peptide 3 as having the highest reactivity with both SARS-CoV-2 Delta wave serums and pre-pandemic DV serums (Fig. 5d). This in vitro assay provided strong in vitro evidence of immunological pressure on these peptide stretches, supporting the hypothesis of cross-reactivity between these two viruses.

From all pieces of evidence, it appears that newer SARS-CoV-2 variants emerged in DV endemic regions under DV and SARS-CoV-2 mutual selection pressures. These culminated in accumulation of increased number of mutations in the RBD and RBM preferably involving residues under positive selection by DV Abs. Whether cross-neutralization potential of DV Abs was diminished for Delta and other variants has not yet been tested by gold standard assays like VNT in cell culture. A lower neutralizing capacity would validate our computational projections.

However, our wet lab experiments have confirmed that our predicted spike residues are indeed, involved in DV Ab binding and prone to sustain mutations to evade DV Ab pressure. Furthermore, experiments by others have demonstrated that SARS-CoV-2 mutations can be triggered by targeted monoclonal antibody therapy, such as casirivimab and imdevimab [50]. The fact that mutated versions reduce or completely abolish an Ab’s affinity and neutralising function suggests that one strong force behind selection is immune evasion. As a result, it is highly likely that DV E Abs, indeed put SARS-CoV-2 and its variants under selection pressure.

Interestingly, we have already reported[10] that with the introduction and evolution of the Omicron variants in India, the previously observed cross-reactivity of SARS-CoV-2 Ab-positive serum samples (mainly against Wuhan and Delta) with DV envelope in DV IgM/IgG detection LFIA test kit decreased considerably. The percentage of DV cross-reactive SARS-CoV-2 serums, collected during January to September 2022 (Omicron waves), was 41.4 %[10]. The cross-reactivity between SARS-CoV-2 and DV was previously noted at 89 % using the same DV IgM/IgG detection assay during the Delta wave [4]. This is in fact, direct experimental evidence that the later variants must have accumulated mutations in the spike to evade binding to DV Abs. That is the reason why Abs to these variants like Omicrons showed almost 50% reduced cross-reactivity in DV IgG/IgM detection assay[10].

The multiple sequence alignment maps of spike amino acid mutations for different variants and respective marked docking regions of DV-2 E Abs indeed, shows a trend of impact of DV Abs pressure (Fig. 4, Table-1). A meta-analysis of a hospital based study suggests that DV co-infection group of patients showed higher rate of mutations in SARS-CoV-2 rather than DV negative group[19]. This report strengthens our predictions; out of 21 amino acid mutations observed jointly in Omicron-BA.1 and BA.2 RBD and RBM regions (this study), 13 showed DV-2 E Abs docking to spike at the same position in case of predecessor variant (s), suggesting that >60% mutations finally observed in the Omicron variants were selected due to change of the original amino acid which showed binding to DV E Abs.

We have also studied the reverse scenario i.e. the impact of COVID-19 in Omicron wave on DV in dengue prone regions like India [10]. Overall, we observed that Abs to earlier strains like Wuhan-hu-1 and Delta (2020-21) were highly cross-reactive in DV serological tests and could neutralize DV1-4 in cell culture [4][10]. This is perhaps one major reason as to why dengue incidences dropped drastically in dengue endemic regions of China, Sri Lanka and Malaysia [11][51].

However, as mentioned before, the Abs to the lately emerged variants like Omicrons were significantly less cross-reactive to DV tests but could still neutralize DV-1, -2 and -4. However, they failed to neutralize DV-3; instead, they caused ADE in case of DV3 in cell culture[10]. This possibly led to an unusual surge of DV3 globally during the Omicron era (2022-23)[10]. This was another example where our experimental results corroborated well with epidemiological observations [4].

Our preliminary analysis showed that SARS-CoV-2 Abs didn’t appear to have perceptible impact on emergence of variants in case of DV serotypes. At least one DV-3 clinical isolate from 2021 showed no change in envelope sequence when compared to that of previous strains isolated in 2019 or even 2017. This suggests that Abs to SARS-CoV-2 and its variants (mainly Wuhan and Delta during 2021) were not robust or persistent enough to impact on DV evolution. However, further studies are warranted before definitive conclusion on this proposition.

Transmissibility of the variants was a different issue-the D614G mutation was shown to augment binding of Delta and other variants to ACE2 leading to greater virus propagation[52][53]. Again, Omicron variants were highly divergent variants and included some of the most critical mutations in the spike protein found to be associated with increased transmissibility and humoral immune escape potential. This has been discussed in detail in our previous study[10]. The impact of DV immunity on the transmission and severity of different variants was variable-Delta caused more infections and deaths; BA.1 caused higher but transient peak of infection than BA.2 which again lingered longer in the population. The impact of DV immunity was also therefore, influenced by the nature of the mutations selected; escape mutations often contributed to increased virulence and/or transmissibility. For instance, one study showed that Omicron-BA.2 was the most infectious variant, which was about 4.2 and 20 times more contagious than the Delta variant and the original SARS-CoV-2, respectively[54].

## 4. Materials and Methods

### 4.1. Modelling and structure refinement of mutated spike proteins

Wild-type SARS-CoV-2 spike protein FASTA sequence was collected from NCBI (MN908947.3). Mutations in the FASTA sequence were generated manually according to reported major mutations characteristic of each variant (Supplementary table 1). Thereafter, I-TASSER server[20] was applied to construct structural models of each mutated spike variant. I-TASSER identified structural templates from the PDB using the LOMETS multiple threading technique, followed by construction of full-length atomic models using iterative template-based fragment assembly simulations. Re-threading the 3D models through the protein function database BioLiP yields insights into the target’s functions. We implemented the I-TASSER as it is a highly reliable server for structure prediction. It has been used for protein structure prediction in recent community-wide CASP7, CASP8, CASP9, CASP10, CASP11, CASP12, CASP13, CASP14, and CASP15 investigations. It was also hailed as the best server for function prediction in CASP9[55]. Standard parameters without steric restrictions were implemented before prediction of the structures. For each spike variant, five models were retrieved. The model with best C-score was selected for further refining using the UCSF Chimera software. All the structures were modified and dock prepared as described previously[8].

### 4.2 In silico docking study of mutated spike proteins with DV-2 E Abs and SARS-CoV-2 Mab in two FFT based global docking serves, HDOCK and Cluspro and image generation in PyMOL

After modifications, for all spike mutated variants, each structure was docked with four DV-2 E Ab (PDB ID: 4UTA, 4UTB, 4UT6, 4UT9) and one SARS-CoV-2 MAb (PDB ID: 7BWJ) in two FFT based global docking servers, namely HDOCK[55] and ClusPro[56] For HDOCK and ClusPro, each antibody structure was uploaded as receptor protein and every spike protein structure was uploaded as ligand protein. In case of ClusPro, special plugins were applied such as masking of non-CDR (Complement Determining Region) for antibody structure. Antibody mode was enabled for docking. Each docked antigen-antibody structure was then analysed in PyMOL using the “find any interaction within 3.5Å cut-off” plugin. Each amino acid of antigens in the interaction interface was identified and marked. The marked amino acids were listed and analyzed in Microsoft Excel. High resolution docked images were prepared by means of the PyMOL Draw/Ray plugin.

### 4.3 Theory and calculation of docking frequency (DF) and cumulative docking frequency (CDF)

For each antigen-antibody pair, total 20 docked structures were analysed. 10 possible docked structures were obtained from each of the servers, HDOCK and Cluspro. First, in an excel sheet, all of the amino acids of RBD that had interacted with one DV-2 E Ab were listed out. Then, the number of times these amino acids interacted with the DV-2 E Ab out of total 20 structures were calculated. The total number of times a DV-2 E Ab interacted with specific RBD amino acids is the docking frequency (DF) of the particular spike protein with one particular DV-2 E Ab. For each variant, the summation of interactions with all 4 DV-2 E Abs was designated as cumulative docking frequency (CDF).

### 4.4 Representative two-dimensional cumulative docking frequency graph and spike protein alignment picture generation

Two-dimensional bar graph of CDF of spike protein variants was created using GraphPad Prism 6 software. Spike gene and amino acid sequences were retrieved from the GenBank and aligned using MEGAX[57] and BioEdit programs.

### 4.5. Data availability

The structures of docking models both in raw and analysed format are available at Mendeley Data (https://data.mendeley.com/datasets/gvvb4rf98h/1). Further information and requests for resources should be directed to and will be fulfilled by the corresponding author, Dr. Subhajit Biswas.

### 4.6. CoV-2 peptide(s) ELISA

Four amino acid stretches in the RBD region were chosen from the SARS-CoV-2 spike WT sequence (NCBI ID MN908947.3). Two dengue virus peptides were also used in the assay as a negative control. NS1Pep1 (N-RRRRRRRRRGKELKCGSGIFKDEL-C) and NS1Pep2 (N-RRRRRRRRRGHTLWSNGVLESKDEL-C) were designed using NCBI sequence of dengue virus 1 NS1 (XDT57230.1). All the peptides were synthesized and coated in 96 well plate using 0.2 M PB buffer (0.2 M phosphate dibasic Na_2_HPO_4_/ NaH_2_PO_4_) in appropriate amount (2.5/10 µg/well). After overnight coating, the wells were washed with washing buffer (0.02 M 1X PBS, pH= 7.2 with 0.05% Tween 20). Wells were blocked with blocking buffer (1% BSA in 0.02 M PBS, pH =7.5). Serum samples were diluted in blocking buffer (1:500) and added to designated wells. After serum incubation, wells were washed with washing buffer and diluted (1:10,000) Rabbit anti-human HRP antibody (ab6759) was added to each well. 3,3′,5,5′-Tetramethylbenzidine (TMB) was added to each well until the colour changed to blue. Reaction was stopped by adding 2N H_2_SO_4_ until the colour changed to yellow. Optical density (OD) of each well was measured using the BioRad iMark^™^ Microplate reader at 450 nm. Signal-to-cutoff (S/CO) ratio was determined for each peptide reactivity with CoV-2 and PPDV serum samples using following equation.

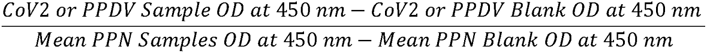

Here, blank OD suggests OD obtained after ELISA for wells without any coating with peptide (s). Raw data of each sample OD can be found in Supplementary table 2. 2D bar graphs of ODs and S/CO ratios were made using GraphPad Prism 6.

## Supporting information

Although SARS-CoV-2 has some proof-reading machinery, numerous mutations began to progressively accumulate in its genome

## Authors’ contribution

S.B. and A.M. conceived and designed the study. A.M. performed all the experiments. All authors analysed and discussed the data. All authors wrote the manuscript to its final version. S.B. supervised the work and critically reviewed and edited the manuscript.

## Acknowledgements

Authors acknowledge the support received from the Director of CSIR_IICB. Subhajit Biswas also acknowledges AcSIR for support. Abinash Mallick acknowledges the support of CSIR for his CSIR Senior Research Fellowship. The research was funded by CSIR_India; grant number: MLP 130 (CSIR Digital Surveillance Vertical for COVID_19 mitigation in India). The grant was given to Subhajit Biswas. The funders had no role in the study design, in the collection, analysis and interpretation of data; in the writing of the manuscript; and in the decision to submit the manuscript for publication.

## Appendix A. Supplementary data

