## Supplementary material for "Role of Dengue in SARS-CoV-2 Evolution in Dengue Endemic Regions": Although SARS-CoV-2 has some proof-reading machinery, numerous mutations began to progressively accumulate in its genome

| SARS-CoV-2 spike variant | Spike protein amino acid mutation |
|---|---|
| B.1.1.7(Kent/UK variant/Alpha) | Δ69/70, Δ144Y, **E484K**, **S494P**, **N501Y**, A570D, D614G, P681H |
| B.1.617 (Delta Variant) | **E484Q**, **L452R**, P681R |
| B.1.617.2.1 (Delta Plus) | T19R, G142D, Δ 156/157, R158G, **K417N**, **L452R**, **T478K**, D614G, P681R, D950N |
| B.1.1529 (Omicron BA.1) | A67V, Δ69-70, T95I, G142D, Δ143-145, Δ211, L212I, Ins.215 EPED, **G339D**, **R346K** **S371L**, **S373P**, **S375F**, **K417N**, **N440K**, **G446S**, **S477N**, **T478K**, **E484A**, **Q493K**, **G496S**, **Q498R**, **N501Y**, **Y505H**, T547K, D614G, H655Y, N679K, P681H, N764K, D796Y, N856K, Q954H, N969K, L981F |
| B.1.1.529 (Omicron BA.2) | T19I, Δ24-27, G142D, V213G, Ins.215 EPED **G339D**, **S371F**, **S373P**, **S375F**, **T376A**, **D405N**, **R408S**, **K417N**, **N440K**, S477N, T478K, **E484A**, **Q493R**, Δ 496- 509, D614G, H655Y, N679K, P681H, N767K, D799Y, Q952H, N972K, |

**Supplementary Table-1**

The table shows key (major) reported amino acid mutations in each SARS-CoV-2 spike variant. Bold mutations denote the ones shown in Fig. 3 and 4. Mutations demarcated in bold letter (point mutations) denote DV-2 E Abs binding sites in predecessor variant which resulted in mutations in Omicron BA.1 and BA.2 variants. Green-highlighted amino acid mutations are the ones which showed DV-2 E Ab(s) binding to a preceding mutant at the same position, which might have resulted in a mutation in Omicron BA.1 and/or BA.2. Yellow highlighted amino acids of Omicron BA.2 are those which have different mutations compared to Omicron BA.1 at the same position under DV-2Ab(s) pressure.

**Supplementary Fig-1**

Multiple sequence alignment of the different successively emerged spike variants (Nucleotide)

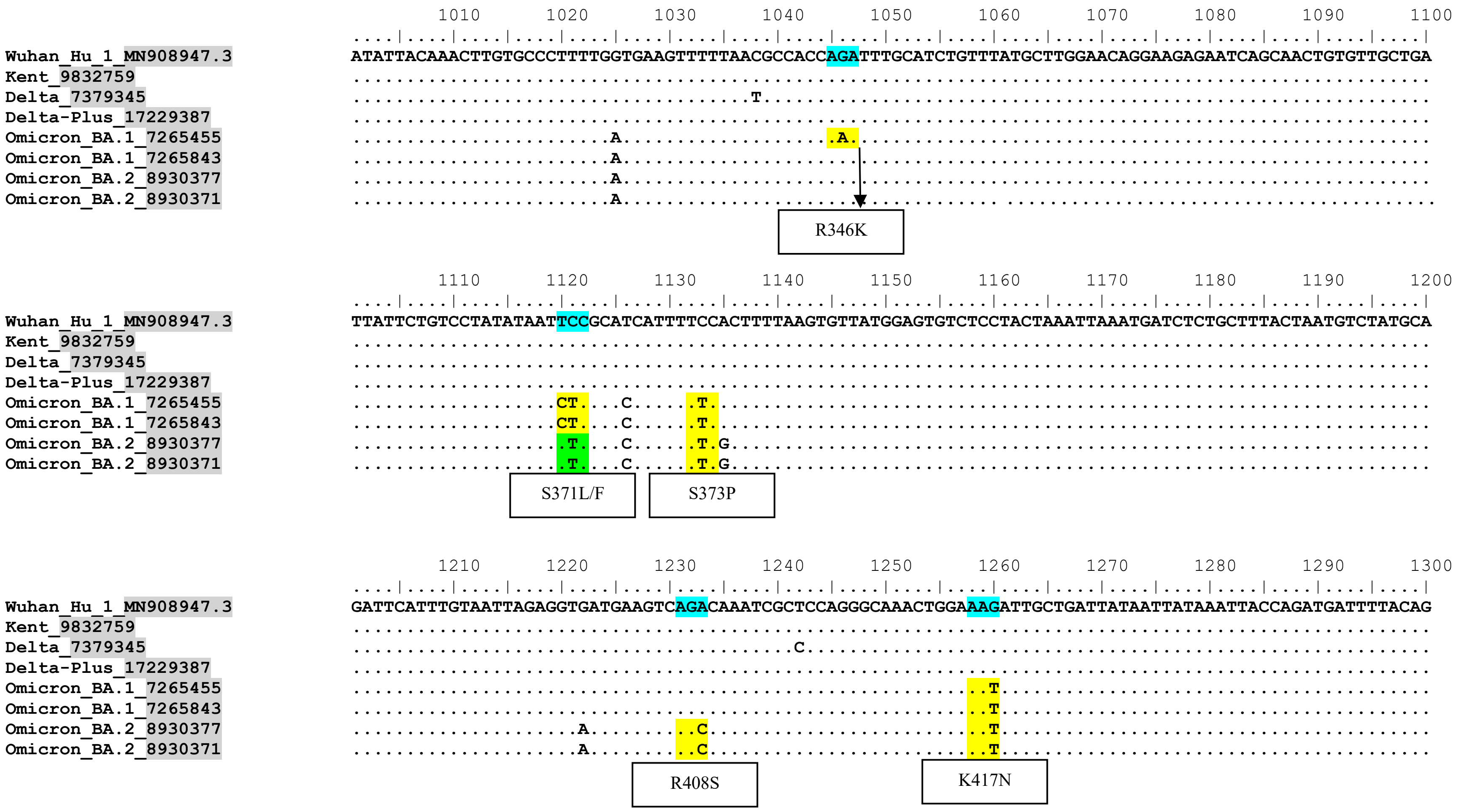

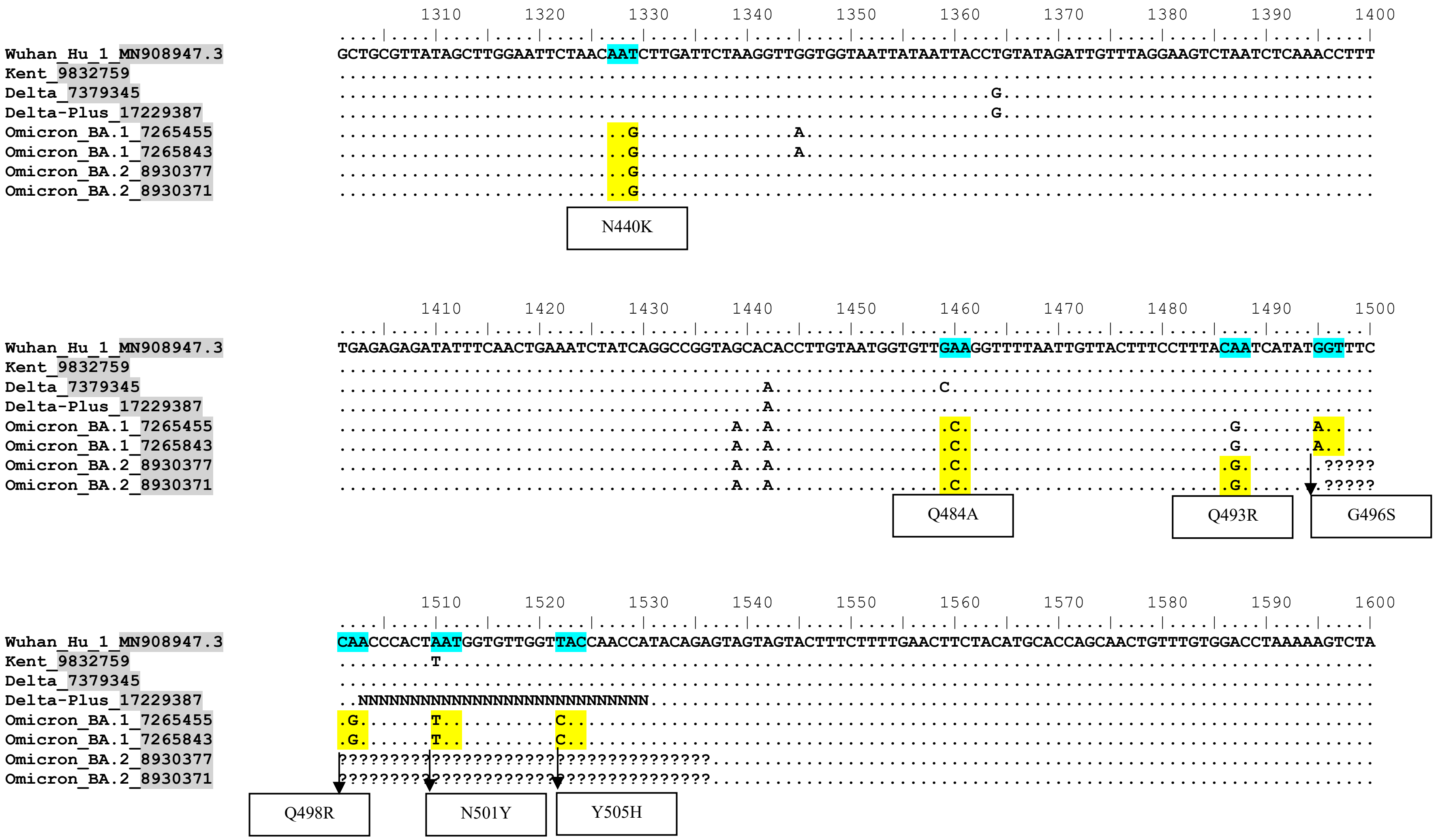
Wuhan_Hu_1_MN908947.3
Kent_9832759
Delta_7379345
Delta-Plus_17229387
Omicron_BA.1_7265455
Omicron_BA.1_7265843
Omicron_BA.2_8930377
Omicron_BA.2_8930371
N440K
Q484A
Q493R
G496S
Q498R
N501Y
Y505H

**Supplementary Table-2**

Raw and analysed data of SARS-CoV-2 synthetic peptide(s) ELISAs

### CoV-2 Peptide 1+2+3+4 (10 µg/well Coating)

| *Sample Name* | *OD 450 nm* | *Blank OD 450 nm* | *S/C Cutoff* |
|---|---|---|---|
| CoV Serum 1 | 2.059 | 0.05 | 4.466429524 |
| CoV Serum 2 | 2.032 | 0.056 | 4.393063584 |
| CoV Serum 3 | 1.978 | 0.068 | 4.246331703 |
| CoV Serum 4 | 1.7 | 0.053 | 3.66162739 |
| CoV Serum 5 | 1.4 | 0.06 | 2.979101823 |
| PPDV 1 | 1.146 | 0.056 | 2.423299244 |
| PPDV 2 | 1.137 | 0.064 | 2.385504669 |
| PPDV 3 | 1.2 | 0.057 | 2.541129391 |
| PPDV 4 | 1.002 | 0.054 | 2.107603379 |
| PPDV 5 | 0.954 | 0.043 | 2.025344598 |

| | *OD 450 nm* | *Blank OD 450 nm* | |
|---|---|---|---|
| PPN 1 | 0.527 | 0.06 | Prepandemic Negative Control Serums |
| PPN 2 | 0.536 | 0.052 | |
| PPN 3 | 0.544 | 0.048 | |
| PPN 4 | 0.532 | 0.07 | |
| PPN 5 | 0.389 | 0.049 | |
| **Average** | 0.5056 | 0.0558 | |

### CoV-2 Peptide 1
### (2.5 µg/well)

| *Sample Name* | *OD 450 nm* | *Blank OD 450 nm* | *S/CO Cutoff* |
|---|---|---|---|
| CoV-2 Serum 1 | 0.875 | 0.05 | 5.449141347 |
| CoV-2 Serum 2 | 0.897 | 0.056 | 5.554821664 |
| CoV-2 Serum 3 | 0.778 | 0.068 | 4.689564069 |
| CoV-2 Serum 4 | 0.743 | 0.053 | 4.557463672 |
| CoV-2 Serum 5 | 0.8 | 0.06 | 4.887714663 |
| PPDV 1 | 0.775 | 0.056 | 4.749009247 |
| PPDV 2 | 0.763 | 0.064 | 4.616908851 |
| PPDV 3 | 0.842 | 0.057 | 5.184940555 |
| PPDV 4 | 0.832 | 0.054 | 5.138705416 |
| PPDV 5 | 0.6 | 0.043 | 3.678996037 |

| | *OD 450 nm* | *Blank OD 450 nm* | |
|---|---|---|---|
| PPN 1 | 0.23 | 0.06 | Pre-pandemic Negative Control Serums |
| PPN 2 | 0.236 | 0.052 | |
| PPN 3 | 0.198 | 0.048 | |
| PPN 4 | 0.133 | 0.07 | |
| PPN 5 | 0.239 | 0.049 | |
| **Average** | 0.2072 | 0.0558 | |

### CoV-2 Peptide 2
### (2.5 µg/well)

| *Sample Name* | *OD 450 nm* | *Blank OD 450 nm* | *S/C Cutoff* |
|---|---|---|---|
| CoV-2 Serum 1 | 0.857 | 0.05 | 8.77173913 |
| CoV-2 Serum 2 | 0.887 | 0.056 | 9.032608696 |
| CoV-2 Serum 3 | 0.8 | 0.068 | 7.956521739 |
| CoV-2 Serum 4 | 0.7 | 0.053 | 7.032608696 |
| CoV-2 Serum 5 | 0.84 | 0.06 | 8.47826087 |
| PPDV 1 | 0.761 | 0.056 | 7.663043478 |
| PPDV 2 | 0.723 | 0.064 | 7.163043478 |
| PPDV 3 | 0.644 | 0.057 | 6.380434783 |
| PPDV 4 | 0.756 | 0.054 | 7.630434783 |
| PPDV 5 | 0.739 | 0.043 | 7.565217391 |

| | *OD 450 nm* | *Blank OD 450 nm* | |
|---|---|---|---|
| PPN 1 | 0.13 | 0.06 | Pre-pandemic Negative Control Serums |
| PPN 2 | 0.139 | 0.052 | |
| PPN 3 | 0.176 | 0.048 | |
| PPN 4 | 0.15 | 0.07 | |
| PPN 5 | 0.144 | 0.049 | |
| **Average** | 0.1478 | 0.0558 | |

### CoV-2 Peptide 3
### (2.5 µg/well)

| ***Sample Name*** | ***OD 450 nm*** | ***Blank OD 450 nm*** | ***S/C Cutoff*** |
|---|---|---|---|
| CoV-2 Serum 1 | 0.984 | 0.05 | 11.04018913 |
| CoV-2 Serum 2 | 1.027 | 0.056 | 11.47754137 |
| CoV-2 Serum 3 | 1.12 | 0.068 | 12.43498818 |
| CoV-2 Serum 4 | 1 | 0.053 | 11.19385343 |
| CoV-2 Serum 5 | 0.9 | 0.06 | 9.929078014 |
| PPDV 1 | 0.816 | 0.056 | 8.983451537 |
| PPDV 2 | 0.863 | 0.064 | 9.444444444 |
| PPDV 3 | 0.744 | 0.057 | 8.120567376 |
| PPDV 4 | 0.754 | 0.054 | 8.274231678 |
| PPDV 5 | 0.7 | 0.043 | 7.765957447 |

| | ***OD 450 nm*** | ***Blank OD 450 nm*** | |
|---|---|---|---|
| PPN 1 | 0.132 | 0.06 | Pre-pandemic Negative Control Serums |
| PPN 2 | 0.139 | 0.052 | |
| PPN 3 | 0.14 | 0.048 | |
| PPN 4 | 0.146 | 0.07 | |
| PPN 5 | 0.145 | 0.049 | |
| **Average** | 0.1404 | 0.0558 | |

### CoV-2 Peptide 4
### (2.5 µg/well)

| *Sample Name* | *OD 450 nm* | *Blank OD 450 nm* | *S/C Cutoff* |
|---|---|---|---|
| CoV-2 Serum 1 | 0.881 | 0.05 | 8.935483871 |
| CoV-2 Serum 2 | 0.866 | 0.056 | 8.709677419 |
| CoV-2 Serum 3 | 0.86 | 0.068 | 8.516129032 |
| CoV-2 Serum 4 | 0.799 | 0.053 | 8.021505376 |
| CoV-2 Serum 5 | 0.812 | 0.06 | 8.086021505 |
| PPDV 1 | 0.74 | 0.056 | 7.35483871 |
| PPDV 2 | 0.696 | 0.064 | 6.795698925 |
| PPDV 3 | 0.805 | 0.057 | 8.043010753 |
| PPDV 4 | 0.742 | 0.054 | 7.397849462 |
| PPDV 5 | 0.77 | 0.043 | 7.817204301 |

| | *OD 450 nm* | *Blank OD 450 nm* | |
|---|---|---|---|
| PPN 1 | 0.136 | 0.06 | Pre-pandemic Negative control Serum |
| PPN 2 | 0.134 | 0.052 | |
| PPN 3 | 0.114 | 0.048 | |
| PPN 4 | 0.195 | 0.07 | |
| PPN 5 | 0.165 | 0.049 | |
| **Average** | 0.1488 | 0.0558 | |
